# Which Index Should I Use? A Comparison of Indices for Enclosure Use Studies

**DOI:** 10.1101/2021.07.04.451046

**Authors:** James E. Brereton, Eduardo J. Fernandez

## Abstract

Enclosure use assessments have gained popularity as one of the tools for animal welfare assessments and Post Occupancy Evaluations. There are now a plethora of studies and enclosure use indices available in published literature, and identification of the most appropriate index for each research question is often challenging. The benefits and limitations of four different enclosure use indices; Original and Modified Spread of Participation Index, Entropy, and Electivity Index were compared. Three artificial data sets were developed to represent the challenges commonly found in animal exhibits, and these indices were applied to these contrived enclosure settings. Three of the indices (Original SPI, Modified SPI, and Entropy) were used to assess a single measure of enclosure use variability. When zones within an exhibit were comparable in size, all three indices performed similarly. However, with less equal zone sizes, Modified SPI outperformed Original SPI and Entropy, suggesting that the Modified formula was more useful for assessing overall enclosure use variability under such conditions. Electivity Index assessed the use of individual zones, rather than the variability of use across the entire exhibit, and therefore could not be compared directly to the other three indices. This index is therefore most valuable for assessing individual resources, especially after exhibit modifications.

## 1. Introduction

Globally, animal collections aim to provide excellent conditions for their inhabitants (Browning & Maple, 2019; Kelling & Gaalema, 2012; Tucker, 2016). However, there remain major challenges with the assessment of welfare. There is no single metric for animal welfare to reliably assess welfare across taxa (Miller et al., 2020), and debate remains as to what the conceptions of good welfare entails (Melfi, 2009). For instance, welfare audits have been developed for commonly housed agricultural species like cattle (*Bos tauros*) (Tucker, 2016), and more exotic species such as elephants (*Loxodonta africana* and *Elephas maximums*) (Carlstead et al., 2013). These welfare audits often include multiple welfare assessment methods. However, many exotic species housed by the zoo and aquarium community, or privately as pets, have yet to be studied in any depth (Melfi, 2009; Rose et al., 2019). Currently, billions of animals of several thousand species are being housed in captivity (Brereton & Brereton, 2020; Mason et al., 2013). Mainstreaming of welfare assessment tools across multiple taxa and industries is therefore valuable (Miller et al., 2020).

Many welfare assessment techniques are now widely available to practitioners. These include qualitative behavioral assessments (Rose et al., 2019) and quantitative behavioral research methodology (Fernandez & Timberlake, 2008; Maple & Perdue, 2013) such as behavioral diversity assessments (Miller et al., 2020), and stereotypy assessments (Kroshko et al., 2016). One tool that has great value across multiple disciplines is the assessment of enclosure use (Ross & Lukas, 2006). Enclosure use has been examined in a range of species housed in laboratories, aquariums, and zoos, using techniques such as Zone Occupancy, Spread of Participation Index (SPI), and Electivity Index measures (for a review, see Brereton, 2020). In addition to their inherent value as welfare audit tools, enclosure use assessments can also be used to measure exhibit value and function as part of a larger Post Occupancy Evaluation (Kelling & Gaalema, 2012).

One assumption made by researchers using enclosure use assessments as a welfare tool is that all exhibit areas (i.e., zones) should have some function (and should therefore be used) by the inhabitants (Plowman, 2003). If zones are avoided by animals, they may be perceived as being unsuitable or of limited value. As a result, many of the commonly used indices measure evenness of enclosure use as a proxy for suitability (Plowman 2003; Prystupczuk et al., 2020). While there are some potential complications to using evenness as a measure of enclosure suitability, these indices should provide a measure as to whether, overall, the animal is using most of the resources made available to it. A carefully designed project can therefore help researchers and practitioners to understand animal’s preferences and develop more biologically relevant enclosures (Rose & Robert, 2013). For example, enrichment might be added to an exhibit, and enclosure use assessment could determine its effects on resource use (Shepherdson et al., 1993). Many types of enclosure use indices are now available to practitioners, so that enclosure use can be measured in a meaningful and repeatable way (Goh et al., 2017; Horikoshi-Beckett & Schulte, 2006; Plowman, 2003).

### 1.1. Application in industry

Care should be applied when assessing the results of enclosure use studies, as the indices seem to suggest that more even enclosure use is desirable (as it implies all enclosure zones are being used well considering their size). However, this is often not true in most captive spaces with differing features that may be unequally preferred. For example, studies on captive greater flamingos (*Phoenicopterus roseus*) revealed that birds used their exhibits unevenly, particularly during the day (Rose et al., 2018). This may be a result of breeding or incubation, or as part of the natural behavior of social animals. The biological relevance of exhibit use should be considered when studies are undertaken: As a colonial bird species, flamingos are likely to be found congregating into dense flocks and as such, uneven enclosure use scores may not be indicative of poor welfare. Conversely, studies on captive European starlings (*Sturnus vulgaris*) in a laboratory setting revealed that the most even enclosure use scores occurred when animals were housed in the smallest cages (Asher et al., 2009). In this case, the smaller exhibits may have been sufficiently restrictive that the birds had limited ability to avoid zones. Uneven enclosure use may in this scenario be indicative of a more appropriate environment.

Some exhibit zones may also be of great value to animals, yet used only for short periods of time. For example, this could be a zone where enrichment devices are available (Clark & Melfi, 2012), high value food resources become available (Troxell-Smith et al., 2016; 2017), or a zone where courtship displays are performed. The comparative value of these zones would not be obvious when using enclosure use assessments. Further tools, such as preference testing, may have merit in these circumstances (Broom, 1988; Fernandez et al., 2004; Fernandez & Timberlake, 2019; Mella et al., 2016).

Whilst most appropriate when used in conjunction with another assessment tool, enclosure use assessments can be particularly valuable when assessing the effect of an exhibit modification (Ross et al., 2009; 2011) or the differences in zone use between individual animals (Clark & Melfi, 2012). For example, studies on the sitatunga (*Tragelaphus spekii*) revealed that there were highly valuable wetland resources in the exhibit, but that some individuals (such as the herd’s immature male) were unable to gain sufficient access to these areas (Rose & Robert, 2014). Identification of zones that are avoided or are competed over may allow animal keepers to make informed decisions about exhibit design, removing exhibit components that are actively avoided.

### 1.2. Types of enclosure use assessment

Not all animal exhibits can be easily carved into equal-sized squares, and as such, several indices and measurements have been developed to aid practitioners. Zones are now often designated based on their biological relevance to animals, and assessments are available for unequal sized zones (Plowman, 2003). For example, exhibits could be divided into zones based on their thermal characteristics, particularly for reptiles and amphibians where this element is critical to exhibit suitability. For research on human-animal interaction, zones could be partitioned based on their proximity to visitors or animal care professionals (Mallapur et al., 2005). Online tools such as Google Earth Pro may be especially valuable when measuring out zones for large, well-established animal exhibits that have existed for considerable time (Horikoshi-Beckett & Schulte, 2006; Rose et al., 2018).

Many enclosure use studies simplify exhibits into a 2-dimensional ‘floor plan’ for the purpose of enclosure use assessment (Shepherdson et al., 1993; Rose & Robert, 2013). For most terrestrial species this is feasible, especially where the animal is unlikely or unable to swim or climb in its exhibit. In aquariums, however, animals may move freely in three dimensions and therefore can access any exhibit area that contains water. For these exhibits, measuring zone use in three dimensions is the most appropriate method (Kistler et al., 2011). However, many rodents and primates climb on the furnishings, walls, and roofs of wired exhibits. For these arboreal species, measuring the exhibit in three dimensions is not necessarily the most appropriate solution. The animal will have access only to those exhibit zones which contain climbing apparatus: it cannot hover in any exhibit area as the aquarium inhabitant might. To solve this issue, Ross et al. (2011) and more recently Browning and Maple (2019) proposed two metrics that allow zone sizes of exhibits to be measured much more accurately. The first of these is conducted by measuring the sizes (e.g., cm^2^) of all useable zones available to the animal, including the equivalent size of each branch or climbing facility available to the animal. The second of these involves measuring the hypothetical space that the animal would take up if it were in these zones (e.g., cm^3^), and then using the dimension to calculate the zone size. Both techniques can be applied to the enclosure use indices described.

In our simulation, we compared four commonly used methods regarding their utility for different enclosure use questions in animal behavior research. These were ‘Original’ and ‘Modified’ Spread of Participation Indices (SPI) (Hedeen, 1982; Plowman, 2003), Entropy (Fernandez & Timberlake, 2019), and Electivity Index (Ross et al., 2011), as described below.

The oldest of the four methods was Original SPI, with its earliest use dating back to the 1950s (Hedeen, 1982). The formula for SPI is:

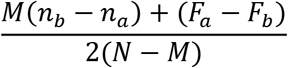

Where N is the total observations in all zones and M is the mean frequency of observations per zone. n_a_ and n_b_ represent the number of zones with observations >M and <M, respectively. F_a_ and F_b_ describe the total observations in all zones with observations <M and >M. The output values for SPI range between 1, indicating selective use of just one zone, and 0, indicating homogenous use of all zones. While Original SPI remains one of the most used enclosure use indices (see Brereton, 2020 for a review), there are several challenges that limit its application to modern exhibits. For example, the index requires all exhibit zones to be of equal size: this is rarely the case in large and complex enclosures (Blowers et al., 2012), especially where irregularly sized dens or branches are available. Even where it is feasible to break an enclosure into equal sized zones, this may be of limited practical value to researchers, as small, valuable resources may be hidden within larger zones (Browning & Maple, 2019).

Considering some of the challenges associated with the Original SPI formula, a second index was developed. Modified SPI was first described by Plowman, in 2003, and was adapted for unequal zone sizes, allowing it to be used for a wider range of animal enclosures containing irregular shapes. The formula for SPI is:

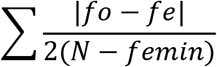

In this formula, *fo* refers to the number of times an animal was observed in a zone, *fe* is the expected frequency for the zone (based on the respective size of each zone), *femin* is the expected frequency in the exhibit’s smallest zone and N refers to the number of observations for the observation period. As per Original SPI, the values for Modified SPI also vary between 1 and 0. This newer index has been used for a range of animal exhibits (Brereton et al., 2021; Rose & Robert, 2013; Rose et al., 2018; Prystupczuk et al., 2019). The ability of the index to assess use on unequal zones has made it increasingly valuable for exhibits containing different substrates and resources (Brereton, 2020). For example, exhibits for amphibious animals such as hippos (*Hippopotamus amphibius*) can be broken down into deep and shallow water and land, so that each resource’s value can be considered.

Entropy is well used in many fields of science to describe changes in structure and has been used to measure enclosure use variability (Fernandez, 2021; Fernandez & Harvey, 2021; Fernandez, Myers, & Hawkes, 2021; Fernandez & Timberlake, 2019). Entropy assesses the amount of randomness or disorder in a system (Fernandez, 2021). The formula for Entropy is:

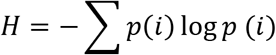

In this formula, *p(i)* refers to the proportion of time spent in the *i_th_* area. This formula produces a number from 0 to 1, in which a value of 1 normally indicates even enclosure use, and 0 is uneven. To improve the comparability of the Original and Modified SPI scores, Entropy scores for this study have been converted to 1-Entropy. Entropy has been used in only a handful of animal-based projects, yet values are much simpler to generate than for either of the SPI formulae. If the two generate similar scores, it possible that Entropy is a good candidate to supersede Original SPI.

All three indices described so far provide a single value for overall enclosure use. The fourth index, Electivity Index, produces use values for each individual zone. Electivity Index was first proposed by Vanderploeg and Scavia (1979) for use in ecology. The formula for Electivity Index is:

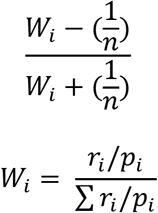

For this index, n refers to the number of zones, r_i_ is the observed use of a zone, and p_i_ is the expected use. The values generated from Electivity Index are per zone and vary from −1 (indicating no use of the zone) to 1 (indicating use solely of that zone). A value of 0 indicated perfect proportional use of a zone in comparison to its size. Much like the Modified SPI formula, Electivity Index overcomes the challenges associated with unequal zone sizes. The index also has some merit in that it can identify over-utilized or under-utilized resources in an exhibit, whereas other indices report only (un)equal enclosure use (Ross et al., 2011). Currently, this index has primarily been used in a handful of primate studies (Goh et al., 2017; Ross et al., 2011). As Electivity Index generates values for each individual resource, it is challenging to compare its use directly to the other indices. However, its functionality, particularly where researchers want to know the value of particular resources to animals, must be stressed.

Considering the range of enclosure use indices available, it is important to understand the benefits and limitations of each technique. In Plowman’s (2003) comparison, blocked zones and differences in frequency of use of those zones was compared for Original and Modified SPI. However, as noted above, for many animal welfare studies, enclosures go beyond flat grids or easily separated zones, and as such, we created three conditions for comparison: (1) small yet valuable resources (similar to that of Plowman’s original comparison), (2) a terrestrial or aquatic habitat, in which exhibit animals can spend time in either land or water, and (3) an aquarium habitat, in which space within the enclosure can be used in a truly three-dimensional manner. In addition, we also included Entropy in the comparisons across all three enclosure types. Finally, we incorporated Electivity Index as a measure for the aquarium habitat to demonstrate its utility in examining differences in valuable resources. We predicted that, regardless of enclosure type, the three compared indices (Original SPI, Modified SPI, and Entropy) would produce similar results when size or value of zones were generally equal. We also predicted that as the differences in size or value between exhibit zones increased, Modified SPI would provide the most sensitive measure to such enclosure use assessments. We discuss all the above with respect to both the validity and applicability of enclosure use indices, as it becomes relevant to consider not only how sensitive any index may be for evaluating enclosure use, but how easy such a metric is to use for the wide variety of animal enclosure types that exist and that we provide a sample of in our simulations.

## 2. Methods

For comparative purposes, artificial data sets were developed to show how each technique might be used. The data sets were simulated so that they could best mimic some of the typical situations and challenges that are faced when researchers investigate captive animal welfare. Three scenarios were generated, consisting of (I) an enclosure containing small, valuable resources, (II) an enclosure containing both land and water, and (III) an aquarium. The contrived data sets were developed for animals living in these exhibits.

### 2.1. Condition I: Small yet valuable resources

For this condition, a basic, rectangular animal exhibit was developed, containing 10 zones of equal size. Each of these zones was further divided into quarters, resulting in forty zones of equal size. Of these forty zones, 10 were set as containing highly valuable resources (in grey) to an animal (see Figure 1).

**Figure 1.**
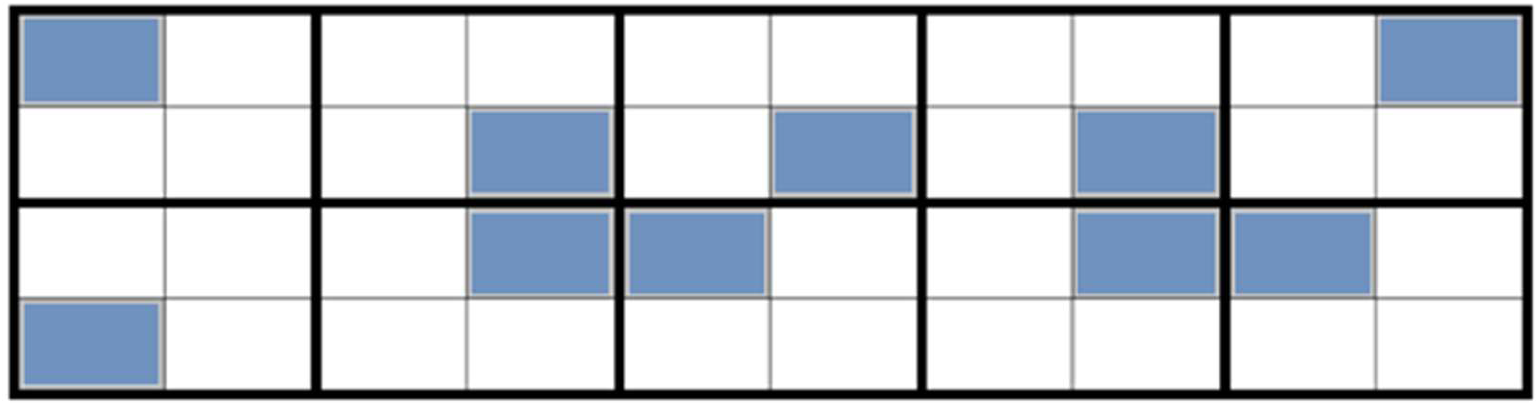
An animal enclosure containing 10 small, desirable resources (grey boxes). The enclosure has been separated into 10 zones (boxes with thick border) and 40 zones (boxes with thin border).

A data set was developed, in which an animal used the valuable resources only and avoided all other areas. The first data point started with the animal using one zone alone, up to the tenth observation where the animal had occupied all ten valuable zones once each. Both Original SPI and Entropy were calculated on this 10-zone (large squares) and Modified SPI was calculated using 11 zones (where each grey box was considered to be a separate zone, and the eleventh zone was the remaining non-valuable exhibit space. Electivity was not used in this comparison as this index generates values for each individual zone.

### 2.2. Condition II: Terrestrial or aquatic habitat

In this condition, a hypothetical animal exhibit was produced containing two biologically different areas, a water feature and a land area. In this scenario, the exhibit was made up of 10 large zones (large black boxes), which were made up of 100 smaller zones in total (small boxes). The land and water took up 75 and 25 zones, respectively (see Figure 2).

**Figure 2.**
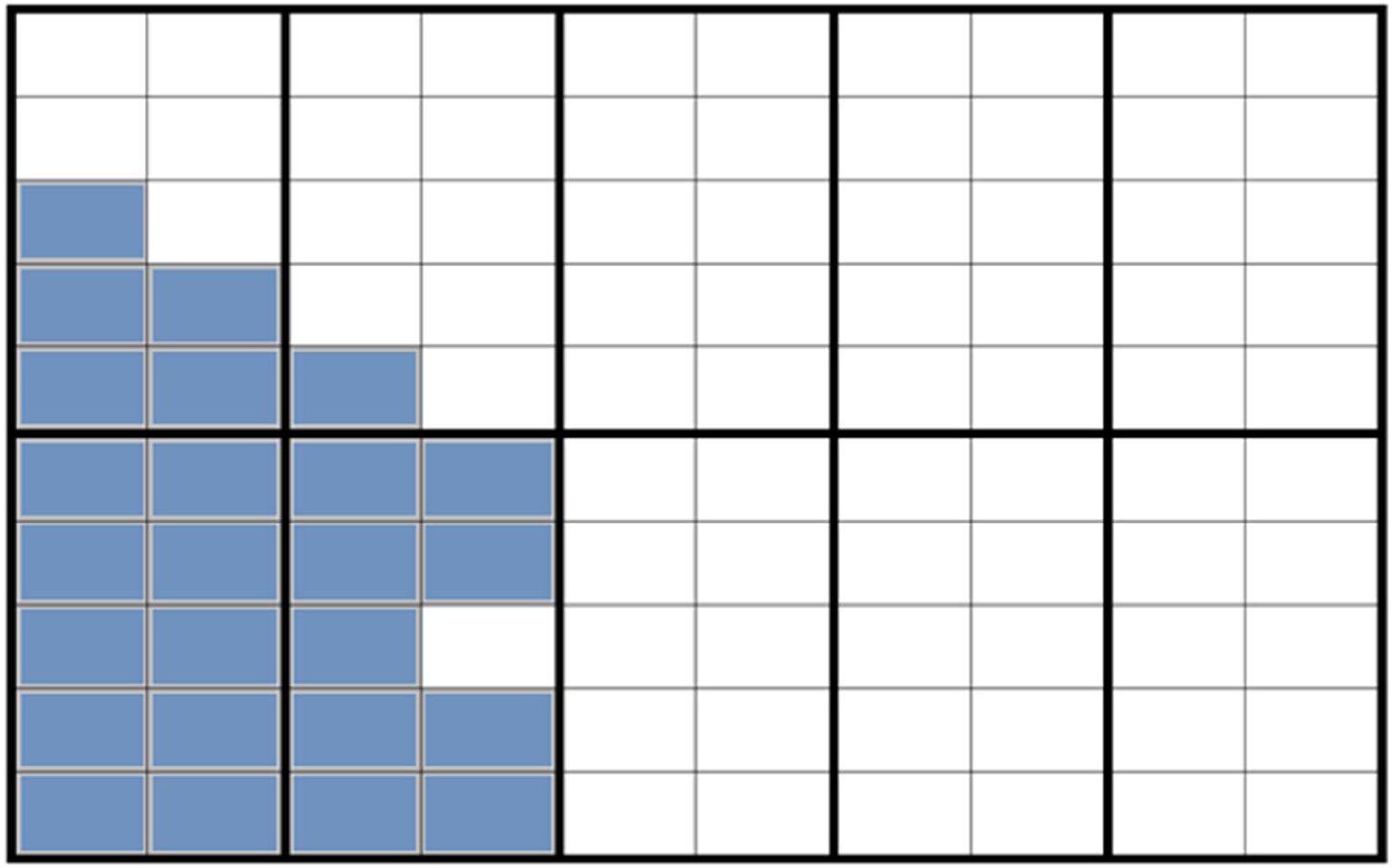
An animal enclosure containing both a water feature (grey boxes) and land (white boxes). The exhibit is divided into 10 equal sized zones for the purpose of enclosure use assessment.

For this study, 11 datasets were developed. In each of these datasets, the animal used 20 zones, each of which was selected using a random number generator. The aim was to determine how an animal’s use of either purely water, purely land, or a mix of both might affect the enclosure use scores. Animals with specializations toward either land or water are found in captivity (e.g., fish), along with animals showing varying degrees of water affinity (e.g., great white pelican (*Pelecanus onocrotalus*)) (Brereton et al., 2021).

For the first dataset, the random number generator was set to select 20 land zones. For the second, the random number generator was set to select 18 land zones (90% of observations) and two water zones (10% of observations). For the third, the number generator was set to 16 land (80%) and 4 water (20%). This pattern was continued (i.e., fourth dataset, 14 land and 6 water) until the eleventh dataset, where 20 water zones (100%) were selected at random. To compare the effect of the four observation types, Original SPI and Entropy were run using an equal-sized ten-zone model. For Modified SPI, the exhibit was divided into two zones based on their biological relevance: water (25 units) and land (75 units). The dataset for the terrestrial-aquatic animal was run using all four indices and the results were compared.

### 2.3. Condition III: Aquarium exhibit

To investigate the application of enclosure use indices for three-dimensional spaces, an aquarium was simulated. The exhibit was divided into four depths, all equal size, for the use of Original SPI and Entropy (see Figure 3).

**Figure 3.**
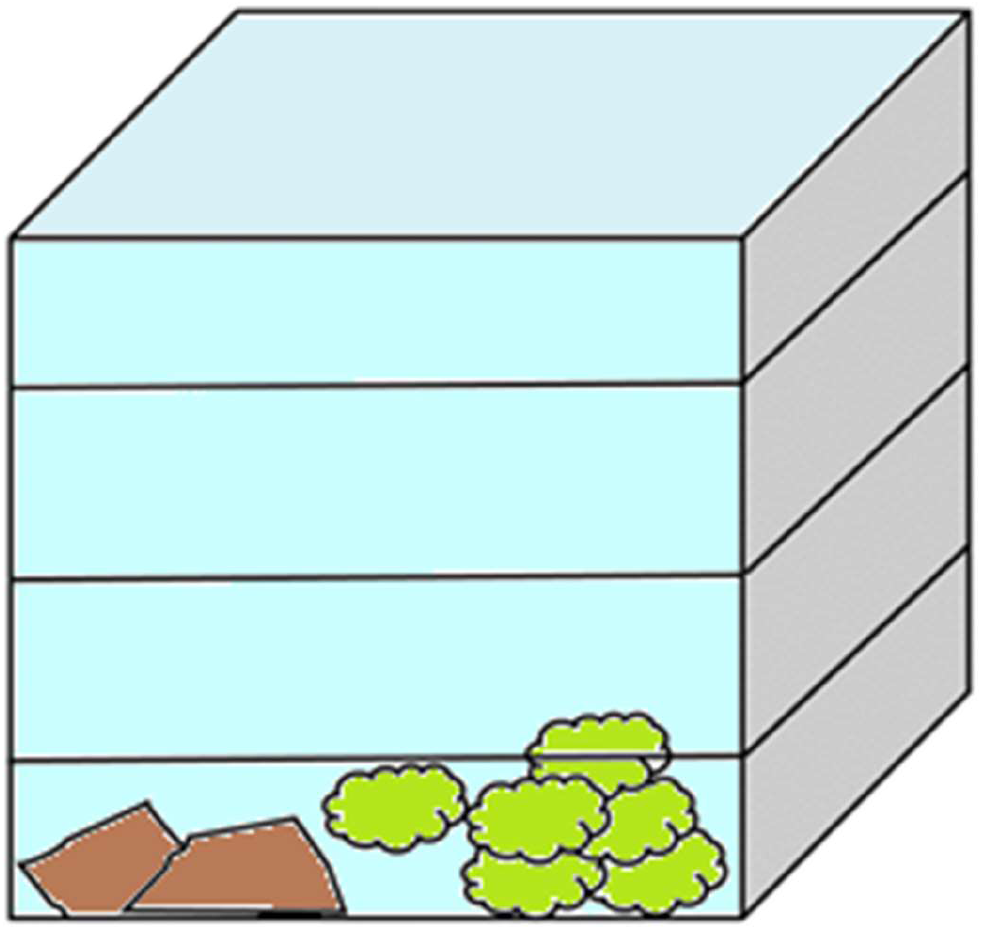
An aquarium exhibit. The exhibit is divided into four equal-sized components which animals have access to.

The enclosure was divided into depths because for many species, water depth is associated with environmental changes (particularly in light intensity, oxygen levels and water pressure). For Modified SPI, the exhibit was divided into four different biological zones: aquatic plants (15%), rocks (10%), the water body (50%) and water’s surface (25%). Electivity Index used the same biological zones as Modified SPI and was covered in a separate figure to the other indices.

A total of 8 simulated datasets were produced. In the first, the animal used all equal-sized zones equally for eight observations. In the second, the animal spent two observations in the rock area (25% of time). For each dataset, the animal spent one more observation in the rock area, until in the eighth, it spent all eight observations in this zone.

## 3. Results

### 3.1. Condition I: Small yet valuable resources

The three enclosure use assessment indices (excluding Electivity Index) were compared (see Figure 4). Overall, both Original SPI and Entropy on 10 zones produced lower scores of evenness of enclosure use, producing values of 0 for the final observation. By contrast, Modified SPI decreased at a steadier pace, showing a much less even assessment of enclosure use.

**Figure 4.**
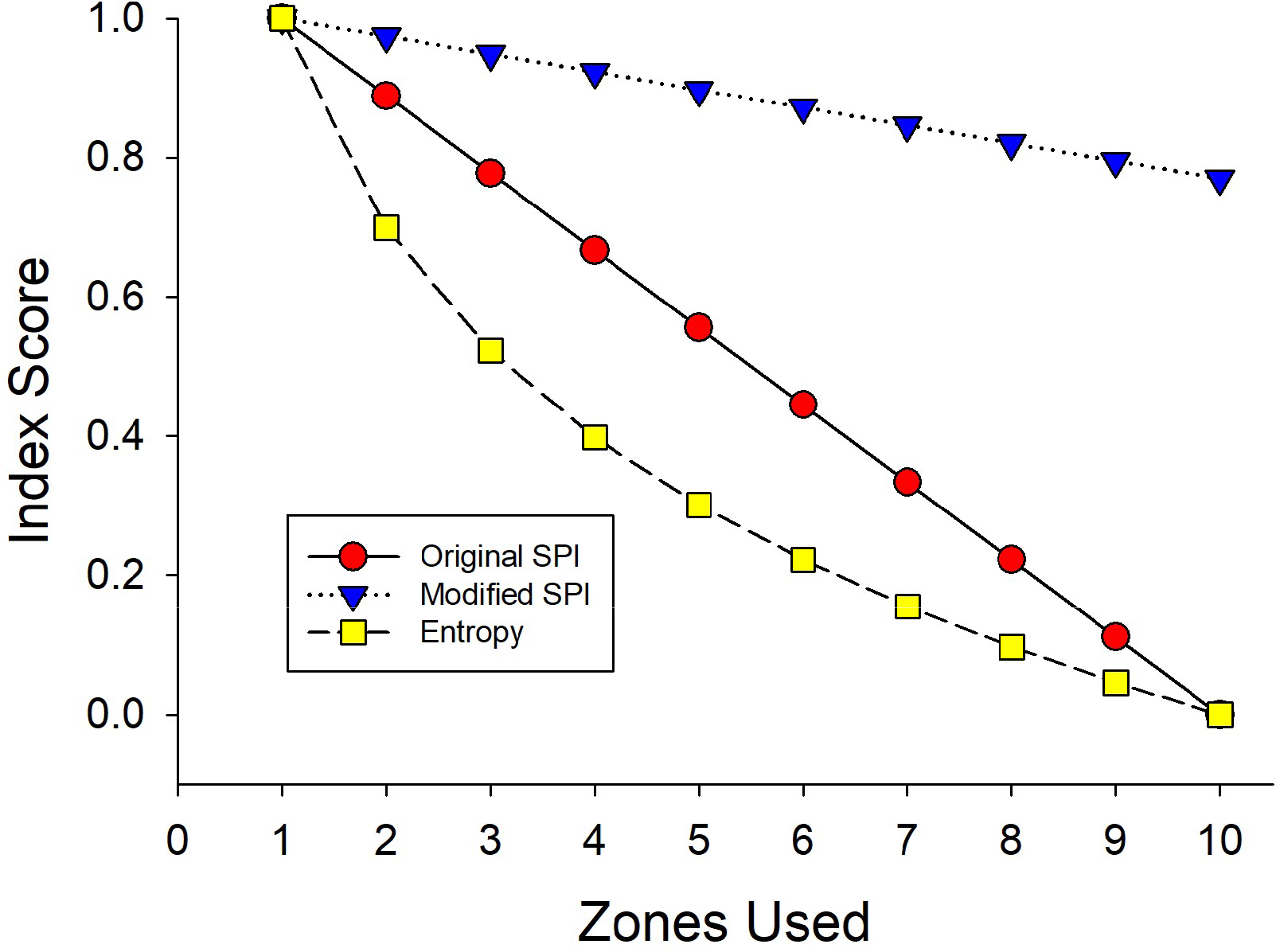
Line chart comparison of Entropy, Original and Modified SPI for a ten observation period.

**Figure 5.**
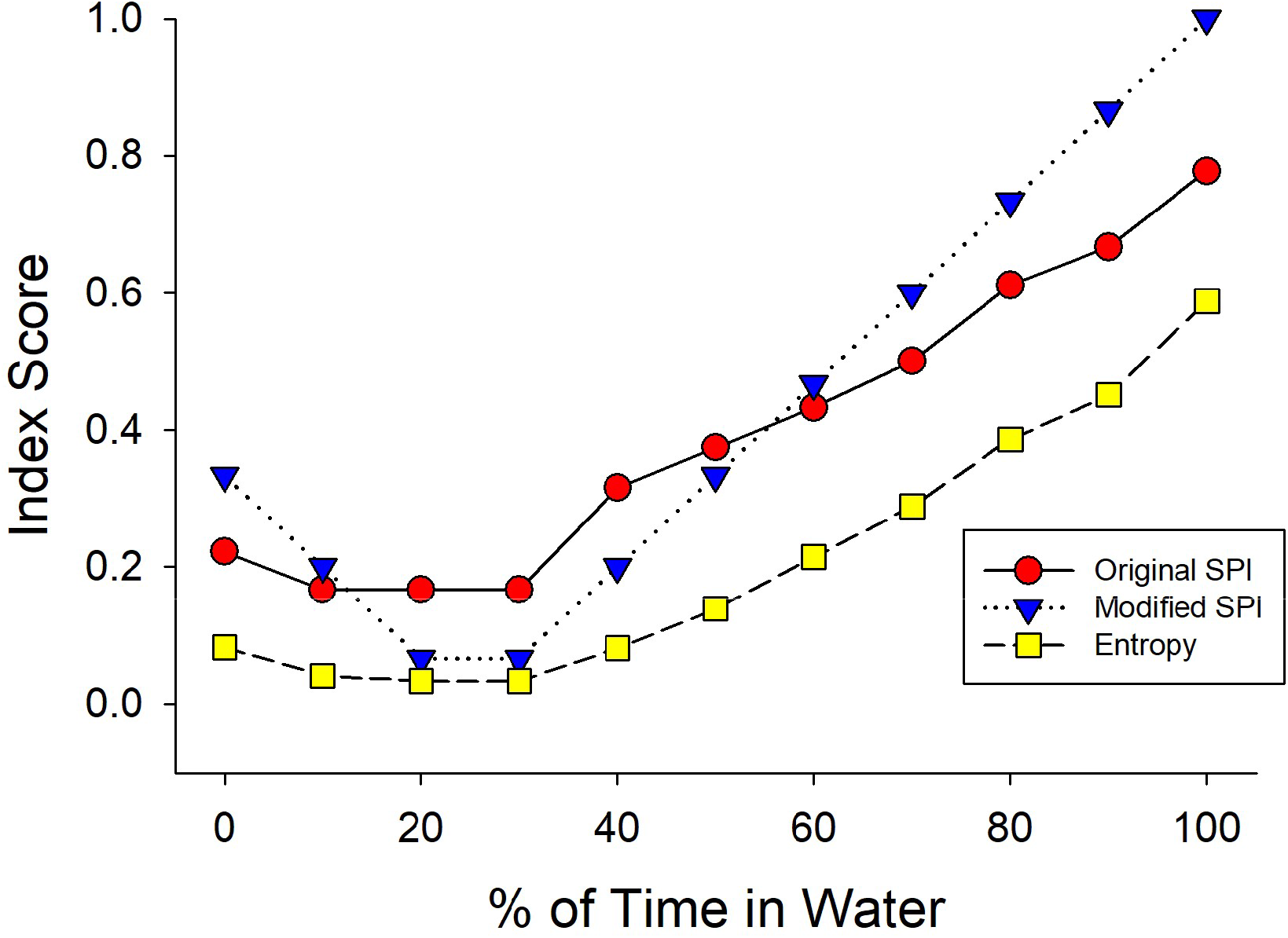
Line chart comparison of Entropy, Original and Modified SPI for an animal with habits ranging from fully aquatic to fully terrestrial.

### 3.2. Condition II: Terrestrial or aquatic habitat

The three indices (excluding Electivity Index) were compared for an exhibit containing water and land, for an animal that changed from entirely aquatic, to entirely terrestrial. None of the indices produced linear lines, with Entropy and both SPI values showing an increase as the species became entirely terrestrial. Generally, Modified SPI scores were highest for the 100% aquatic animal, and the lowest score was for Entropy.

### 3.3. Condition III: Aquarium exhibit

The indices were compared for an aquatic enclosure containing four Original, equal-sized zones, and four biological zones (water surface, water body, rocks, and aquatic plants). Figures 6 shows the index scores as the animal transitions from using all zones equally, to using only the rocky zone. The comparison of the Original and Modified SPI, and Entropy indices show that all indices are in agreement when the most uneven enclosure use scores are shown. Entropy shows the highest values when the animals are using their exhibits relatively evenly.

**Figure 6.**
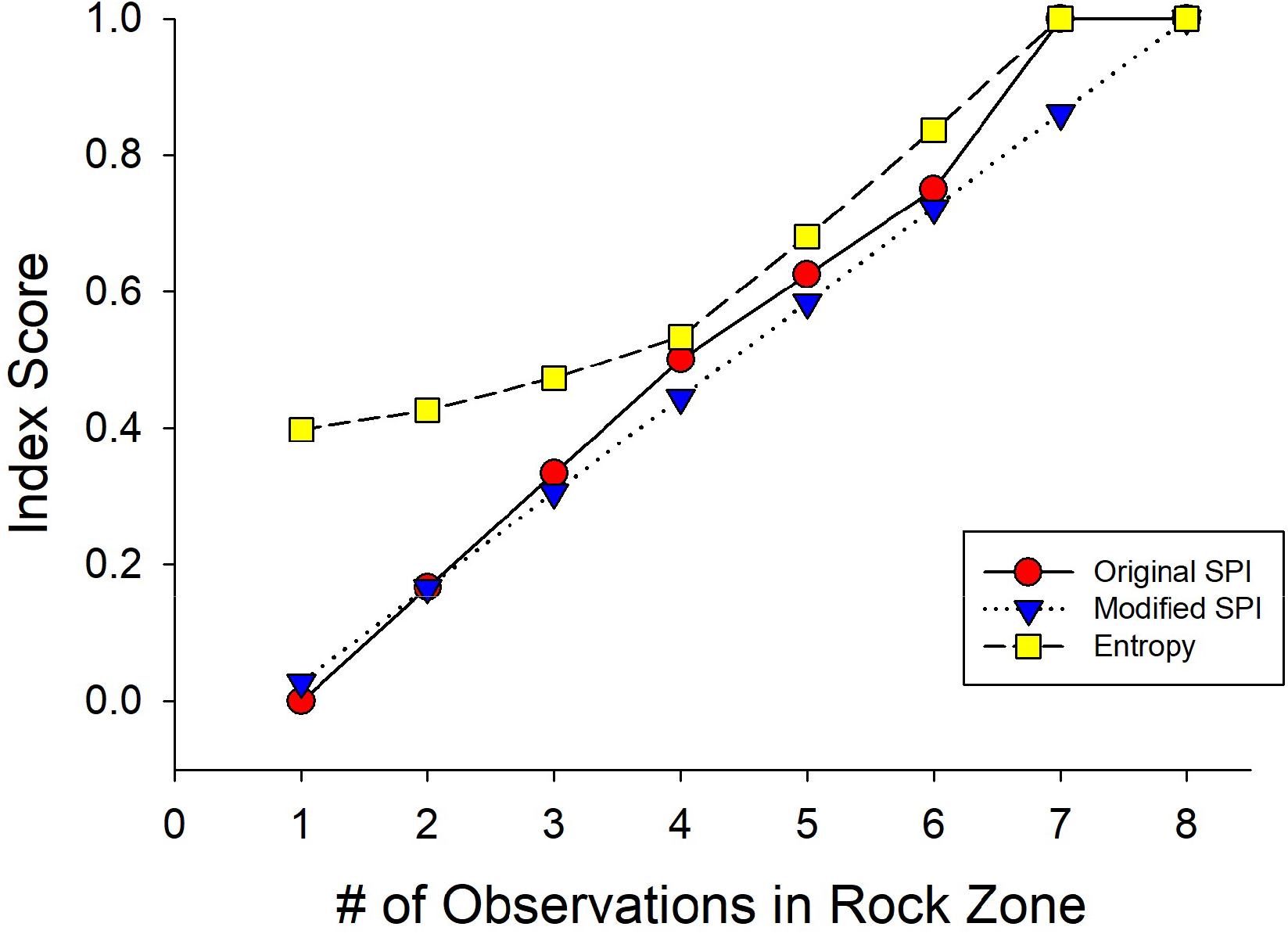
Line chart comparison of Entropy, Original and Modified SPI for an aquatic animal, which uses its exhibit evenly (observation 1), through to an animal that shows preference for just the rock zone (observation 8).

Electivity Index (see Figure 7) was run separately to the other indices, using the same data on aquarium use. The Index shows a breakdown of zone use change and identifies whether the zone is being over or under-utilized in comparison to its size.

**Figure 7.**
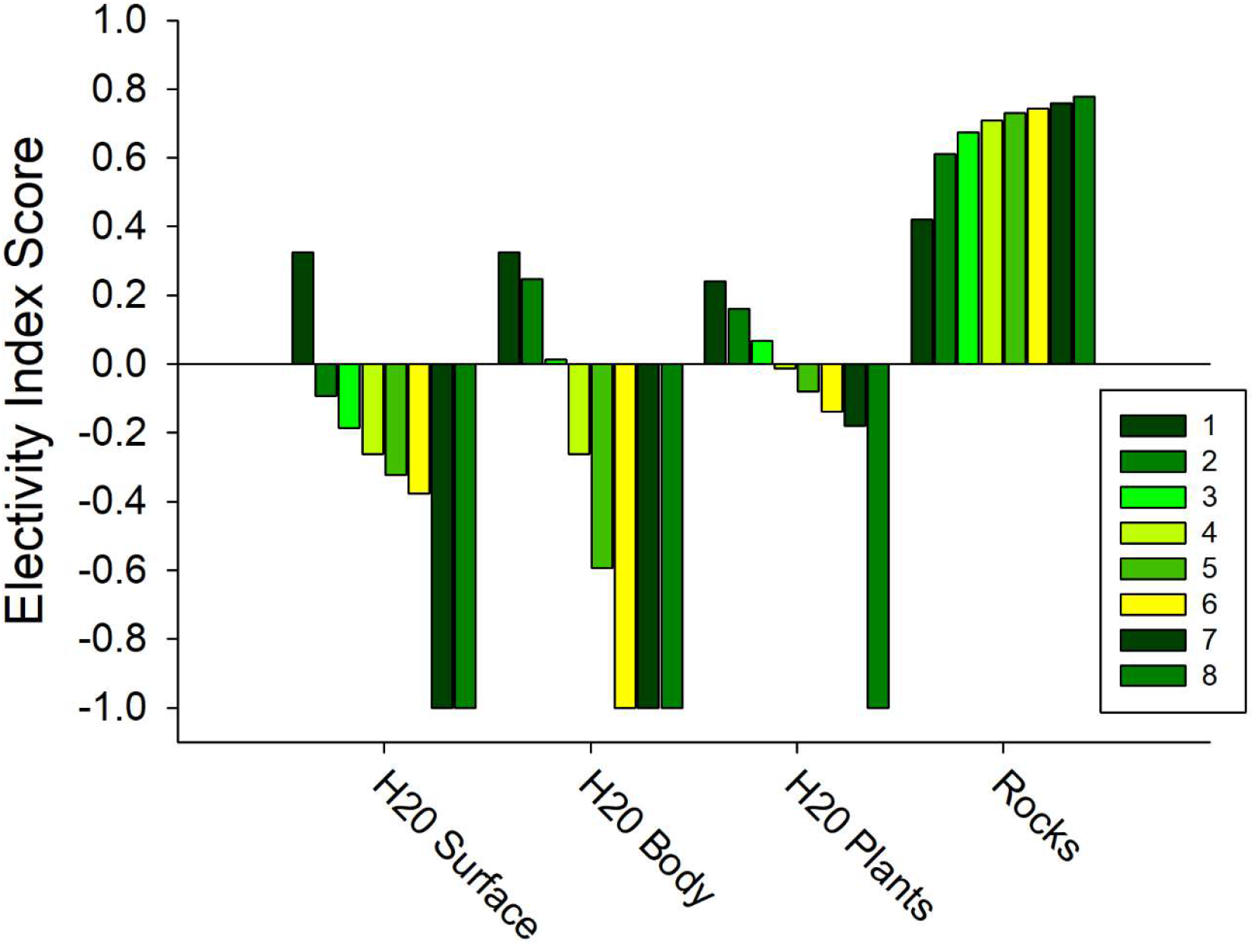
Electivity scores for an animal that initially uses its aquatic enclosure evenly, through to an animal that shows preference for just the rock zone. A score of 1 and −1 indicate exclusive use and non-use of a zone respectively. The key from 1-8 indicates the number of observations used to generate the data.

## 4. Discussion

We were able to effectively compare three of the indices (Original and Modified SPI and Entropy) across three different conditions: an exhibit with small, valuable resources, with both land and water space, and an aquarium with four equal zones of different depths. We also showed how Electivity Index is used for the 3-dimensional exhibit. Our results can be summarized with the following three points: (1) The indices used produced similar results the more equally each zone was used, (2) Modified SPI was most effective for assessing enclosure variability when zones were less equally utilized, and (3) Electivity Index was effective at measuring substrate/zone preferences, but inappropriate for measuring overall enclosure variability.

### 4.1. Condition I: Small yet valuable resources

The three comparative indices; Original and Modified SPI and Entropy all clearly showed scores closer to 0 when animals used their enclosures more evenly. For the first simulation, on small yet valuable resources, Original and Modified SPI scores decreased in a linear fashion as enclosure use became more even. Entropy, by contrast, which has been used less frequently in enclosure use research, did not (Fernandez & Timberlake, 2019). The Entropy scores dropped more steeply than either Modified SPI, or Original SPI, with which it used the same zone measurements.

In Condition I, Modified SPI was able to incorporate a much more detailed view of the animal’s enclosure, and therefore its use of the exhibit. For example, in the simulation where the animal was using all ten small valuable resources but no other areas, Modified SPI scores remained high. By contrast, both the original SPI and Entropy scores had dropped to 0. According to both original SPI and Entropy, the animal was technically using all zones equally, so the score of 0 is not surprising. Modified SPI, by contrast, was able to more appropriately identify that the animal was using 25% (10 of 40 blocks) of its exhibit.

The strength of Modified SPI, therefore, is in providing the researcher with the ability to divide their enclosure into zones that best reflect the nature of their research question. For a project relating to visitor effects, the researcher may assign zones based on their proximity to visitor pathways, or based on biological relevance (Prystupczuk et al., 2020). The SPI values produced would therefore assess the evenness of enclosure use in consideration of visitors, or functional value of the environment.

Entropy, which is a measure of variability, produced generally similar results compared to the former two indices under the small, valuable resources simulation. The Entropy values were slightly lower, but similar to the values demonstrated by Original SPI. However, the difficulty in interpreting such results is that (a) most exhibits consist of space that is seldom uniform, and (b) animals rarely utilize exhibit space in an unbiased or randomized, equally distributed fashion (Fernandez & Timberlake, 2019; Kelling & Gaalema, 2011). Therefore, one clear suggestion for examining overall enclosure use would be to create zones animals can use that are both equal in size as well as in expected frequency used. Care should be taken when dividing any enclosure space into separate zones in attending not only to the differences between each zone (i.e., water vs land zones), but in the actual size and perceived frequency of usage of the designated zones.

### 4.3. Condition II: Terrestrial or aquatic habitat

The purpose of this simulation was to demonstrate how each of the indices performed on an exhibit containing both land and water. In this exhibit, the land and water took up 75% and 25% of the available space, respectively. Intuitively, one might expect the lowest index scores when the animal spent 20-30% of its time in the water: this was indeed the case, as both Modified SPI and Entropy scores were close to 0 for these observations. The Original SPI score, by contrast, was higher than the other two indices. This is likely due to Original SPI’s limited ability to focus on specific enclosure features: it could not accurately assess the use of water, as water was concentrated within only a few zones. It is surprising, then, to see that Entropy, which used the same equal-sized zones as Original SPI, showed greater similarity with Modified SPI scores when the animal spent 20-30% of its time in water. The similarity between Entropy and Modified SPI became less apparent as the animal used its enclosure less evenly (up to 100% of its time in water). Here, Modified SPI produced much higher values than either Entropy or original SPI: this is once again likely a result of the differences in enclosure separation.

### 4.4. Condition III: Aquarium exhibit

The aquarium simulation was used to show researchers how each index might fare in an exhibit where measuring each zone in cubic rather than squared measurements might be more appropriate. In this condition, the scores for Original and Modified SPI were much more similar to each other than in previous conditions. Entropy scores appeared higher than the other indices. Electivity Index was also used, though not compared to the others, since Electivity does not produce a similar measure of enclosure variability (Goh et al., 2017).

In this condition, either Original or Modified SPI may be equally appropriate, depending on the study question being asked. In a study where depth is of interest, Original SPI may be as appropriate at Modified SPI, especially in a traditional rectangular aquarium which can be easily divided into equal sized depths. However, it should also be noted that some aquatic species will selectively occupy specific depths (e.g., Kistler et al., 2011); in this case, depth becomes a biologically important characteristic.

Electivity Index was effective at measuring preferences for variables related to different zones, such as substrates or fixtures within a zone. However, the index was associated with challenges at assessing the suitability of an enclosure holistically. For example, the index was not appropriate for the first of the contrived data sets. Electivity, whilst unsuitable as an overall measure, excelled in showing the under- and over-utilization of different resources in relation to their size (Goh et al., 2017). Electivity may therefore be particularly appropriate for projects where one zone (containing a new resource or enrichment features) is of interest.

### 4.5. Future Directions for Enclosure use Indices

Early studies divided exhibits into rectangular, equal-sized zones to assess enclosure use (Hedeen, 1982), with a greater diversity of studies developing once non-equal zone indices became available (Plowman, 2003; Rose & Robert, 2013; Ross et al., 2009; 2011). With the expanding remit of welfare assessments, enclosure use studies may now be able to answer a great diversity of questions. For example, future research could use a mix of behaviour and enclosure use assessment to develop a more informed picture as to how exhibits are used. Using a technique such as Electivity could identify which zones are overutilized, and behavioural research could then identify what behaviours the animals conduct when in each of these zones. This could allow researchers to determine the potential value of specific zones, especially where they are rarely in use (Maple & Perdue, 2013).

From the literature, it appears that enclosure use assessments are not used uniformly in all animal sectors (Brereton, 2020). For example, there is good evidence to show enclosure use is present in welfare audits in laboratories (Asher et al., 2005), aquariums (Kistler et al., 2011) and zoos (Brereton, 2020). However, there appears to be limited published information on enclosure use in the farm setting. Here, enclosure use measurement could have merit in assessing the value of different resources, allowing farmers to make informed decisions about resource provision and exhibit design. Likewise, technology has been developing rapidly, and there is some synergy for its use in enclosure use studies. For example, several studies have made use of satellite technology to measure zone sizes for large, open-air exhibits (Rose et al., 2018; Rose & Robert, 2013). For other studies, Global Positioning Service (GPS) assessments have been used to identify animal locations (Blowers et al., 2012; Leighty et al., 2010). Trackers for animals, or automated recording techniques, could be used to reduce the requirements for human observers, or observe animals during closed hours (Rose et al., 2018). Specifically, nocturnal studies could provide more insight into the relevance of exhibit resources, particularly for species that are typically crepuscular or nocturnal.

## 5. Conclusions

Based on our simulations and the nature of using enclosure use indices, we conclude the following: (1) Enclosure use indices should be used in conjunction with other welfare measures, as well as determined through both individual and species-typical exhibit use considerations. When used without other assessments, enclosure use has limited value in assessing animal welfare. (2) When possible, enclosure use assessments should split exhibits into zones of both comparable size and value. The more equal the zone size or value, the easier it is to evaluate the impact of changes on such zone usage on the welfare of an animal under different conditions. (3) When zones are equal in size or value, Entropy, Original SPI, and Modified SPI are near equally effective at assessing overall enclosure use. As zones diverge from equal zone size or value, Modified SPI becomes a more sensitive measure for assessing such conditions. Additionally, Electivity Index is most useful in assessing the importance of specific zones or resources. (4) Finally, there is necessarily a tradeoff between the sensitivity and complexity of any enclosure use index. For instance, Entropy is one of the easiest indices to calculate and under all simulations was effectively able to identify changes in overall enclosure variability. As such, while Modified SPI may function as a more sensitive metric, Entropy may still be more pragmatic for a variety of welfare assessment needs. Welfare assessments should be guided by both the validity and applicability of the tools used. It is our hope that this paper makes the evaluation of enclosure use indices a more viable tool for all welfare scientists.

## Author Note

The authors are grateful to Mrs. S Brereton for proof-reading the manuscript. The authors would also like to thank two anonymous reviewers for providing constructive feedback to help improve the previous version of this manuscript.

This work received no funding, and we have no known conflicts of interest to disclose.

